# The C-terminal region of interferon-γ mediates evolutionarily tunable membrane interaction

**DOI:** 10.64898/2026.11.13.766334

**Authors:** Katarina Baxová, Šárka Pokorná, Hana Rozhoňová, Adam Hruška, Aditi Konar, Bailasan Haidar, Jarmila Sekerešová Králová, Jiří Zahradník

## Abstract

Interferon-γ (IFN-γ) is a key cytokine orchestrating innate and adaptive immunity, yet the functional significance of its highly basic C-terminal region remains poorly understood. Here, we combine evolutionary, biophysical, cellular, and computational approaches to uncover the membrane-interacting properties of this region across vertebrate IFN-γ orthologs. We show that the previously described conserved KRKR motif is embedded within a broader cationic region that is highly diverse in fish but progressively reduced in terrestrial tetrapods. Molecular dynamics simulations and experiments using large unilamellar vesicles reveal preferential interactions with anionic membranes. Furthermore, all tested IFN-γ-derived peptides associate with U-2 OS cells, with the olive flounder-derived peptide displaying exceptionally strong cellular association and cell penetration, markedly exceeding that of the canonical cell-penetrating peptide R_9_. Together, our findings identify the IFN-γ C-terminus as an evolutionarily diverse, membrane-interacting module with pronounced cell-penetrating properties in certain fish orthologs, revealing an unexpected functional dimension of this conserved immune protein.

## Introduction

Interferon gamma (IFN-γ), also classified as type II interferon, was initially identified as a mediator of cellular resistance to viruses. Later studies showed that it is one of the key players in the immune system of vertebrates, bridging innate and adaptive immunity [1]. Its functions are truly pleiotropic, orchestrating a broad range of responses including antiviral defence, antibacterial defence, immunomodulation, and anticancer activities by inducing interferon-stimulated genes (ISGs) [2, 3]. As the most potent cytokine in macrophage activation and promotion of CD4+ differentiation and CD8+ cytotoxic T cell responses, IFN-γ gained a position as a signature Th1 cytokine (proinflammatory response cytokine) [4, 5].

IFN-γ emerged early in the evolution of jawed vertebrates (Gnathostomes), alongside the ancestral members of the interleukin-10 family [6], and subsequently diverged rapidly [7]. Its presence is well documented in extant cartilaginous fishes, bony fishes, and all tetrapods. In higher vertebrates, IFN-γ is represented by a single gene. However, additional type II interferons are present in many bony fish species, including IFN-γ-related (IFN-γrel) proteins. In some cases, multiple copies of these genes exist due to whole-genome duplication events [8].

Human IFN-γ is secreted as a noncovalent homodimer with an intertwined six-helix bundle structure [9]. Each subunit of the mature IFN-γ homodimer is glycosylated at a conserved N-linked site, which, while not essential for biological activity, serves to protect the cytokine from proteolytic degradation [1]. IFN-γ signals through a heterotetrameric receptor complex consisting of two interferon gamma receptor 1 (IFNγR1) chains and two interferon gamma receptor 2 (IFNγR2) chains. The structure of the IFNγ receptor complex was solved using an affinity-enhanced IFNγR2 variant, which inherently binds with lower affinity than IFNγR1 and acts as the primary modulator of signal transduction (PDB 6E3K, [2]). Each receptor chain belongs to the class II cytokine receptor family and features two fibronectin type III domains in the extracellular region, while the intracellular domains are intrinsically disordered [10, 11].

IFN-γ signals through the Janus kinase/signal transducer and activator of transcription (JAK/STAT) pathway. This pathway involves three key components: cytokine receptors, Janus kinases (JAKs), and signal transducer and activator of transcription proteins (STATs). Upon IFN-γ binding to its heterodimeric receptor (IFNγR1/IFNγR2) on the surface of target cells, the associated kinases JAK1 (bound to IFNγR1) and JAK2 (bound to IFNγR2) are brought into proximity. These kinases transphosphorylate and activate one another, subsequently phosphorylating STAT1 on Tyr701. Phosphorylated STAT1 then forms homodimers known as γ-activated factor (GAF), which translocate to the nucleus and bind to gamma-activated sequence (GAS) elements to initiate transcription of ISGs [3].

However, this textbook cascade oversimplifies the regulatory complexity of IFN-γ signalling. First, Zoler et al. demonstrated that multiple JAKs can bind competitively to interferon-α (Type I IFN) receptors depending on their relative abundance, and that such promiscuous JAK recruitment is a general feature shared by type I and II cytokine receptors [12]. Second, STAT1 can form heterodimers with STAT2, establishing a strong functional crosstalk between type II and type I IFN signalling pathways [13, 14]. Additional heterodimers between STAT1 and other STAT family members likely mediate broader signalling interactions with other cytokines, particularly those in the IL-10 family [15].

Third, while the classical understanding of IFN-γ signalling has emphasised cell surface receptor binding and downstream STAT1 activation, emerging evidence suggests that the C-terminal region of IFN-γ also plays a crucial, evolutionarily conserved role in regulating its biological activity. As early as the late 1980s, it was shown that the C-terminal segment of IFN-γ (approximately residues 80–143) is essential for full biological activity, and that its truncation leads to a substantial loss of function [16–18]. This C-terminal region contains a polybasic motif with a canonical nuclear localisation signal (NLS). This sequence is important not only for the efficient nuclear import of IFN-γ itself but also for promoting the nuclear translocation of STAT1α and, potentially, the IFNγR1 receptor subunit. Notably, deletion of residues 95–133, encompassing this polybasic region, prevents STAT1 activation despite preserved receptor binding [19]. Unfortunately, none of the currently available structural models of IFN-γ orthologues and their receptors show a resolved interaction between the polybasic C-terminus and the receptor. Although some studies claim full signalling capacity mediated by the C-terminus only [20], more recent studies have further questioned this mechanism by showing that IFN-γ variants lacking the terminal KRKR motif are still capable of binding to IFNγR1 and retain biological activity [21]. It is therefore evident that important biological functions of the IFN-γ C-terminus remain incompletely understood and require further investigation.

An additional function of the C-terminal region of IFN-γ is to promote its local retention at sites of infection or inflammation, thereby limiting systemic dissemination and associated toxicity. Systemically elevated interferon-γ (IFN-γ) levels, particularly when sustained over prolonged periods, are associated with adverse physiological effects [22–24]. Previous studies have demonstrated that the whole protein associates with negatively charged liposomes [25] and that the C-terminal region of IFN-γ interacts with heparan sulfate in the extracellular matrix, facilitating its retention at the site of secretion, thereby substantially limiting its systemic dissemination and mitigating toxicity [21, 26].

In this study, we show that the C-terminal domains of IFN-γ orthologues exhibit strong affinity for biological membranes. Furthermore, in *Paralichthys olivaceus* (olive flounder) and potentially other cases, C-terminal domains of IFN-γ display passive cell-penetrating peptide-like (CPP-like) properties similar to oligoarginines. These findings further support a conserved functional role for the IFN-γ C-terminus in mediating spatial restriction of cytokine activity and highlight its broader relevance in modulating immune signalling.

## Results and discussion

### IFN-γ proteins contain a conserved and diverse positively charged C-terminal motif architecture

In our previous study investigating type II interferons in teleost fish, we observed striking differences between the C-terminal regions of canonical fish IFN-γ and a group of related proteins termed IFN-γ–related (IFN-γrel) proteins [8]. Notably, IFN-γrel proteins exhibited considerable variation at the C-terminus, with many sequences being markedly truncated or entirely lacking this region. The experimental structure of the IFN-γrel protein from *Ctenopharyngodon idella* [27] with a truncated C-terminal region further confirmed this distinction and preservation of overall IFN-γ architecture. While both IFN-γ and IFN-γrel proteins have been shown to exhibit antiviral activity via STAT1-dependent signalling pathways [28], the functional divergence between these paralogues remains understudied. It has been hypothesised that they mediate distinct immunological roles and the unusual C-terminal region likely plays a role [10]. In particular, the canonical IFN-γ protein from *P. olivaceus* (PoliIFN-γ) possesses a C-terminal sequence characterised by eight consecutive arginine residues, accompanied further by five additional positively charged amino acids (sequence NTKMQRRRRRRRRQARKVKTPTRA). Surprisingly, BLASTP analysis of this arginine-rich C-terminal sequence revealed significant similarity not only to other fish interferons but also to proteins of bacterial origin containing analogous poly-arginine motifs (e.g., Sequence ID: MCB9637973.1), as well as viral proteins, including HIV-1 Rev protein (e.g., Sequence IDs: UHM26946.1, AAK15482.1). A review of the literature on arginine-rich sequences highlighted their frequent association with CPPs [29, 30] or multimerisation [31, 32]. These observations prompted us to hypothesise that the IFN-γ C-terminus may facilitate targeted interactions with biological membranes via CPP-like properties.

To explore the diversity of IFNγ proteins and the distribution of potential CPP-like motifs across all chordates, we compiled a comprehensive dataset of IFN-γ homolog protein sequences by combining NCBI Protein database queries with iterative BLASTP searches restricted to the Chordata taxon. Our search retrieved a dataset of 342 bony fish and 491 non-fish chordate IFNγ sequences. To identify potential CPP-like motifs, we scanned the C-terminal 20 amino acids of each IFNγ homolog for exact matches to 1564 known L-chirality CPP motifs from the CPPsite 2.0 database [33]. In cases of multiple matches, the longest motif was retained. Finally, to assess phylogenetic relationships within selected chordate groups, we constructed maximum-likelihood trees using trimmed protein alignments and robust bootstrap support. These trees are shown in Fig. 1A, B, and in Supp. Fig. S1, including the presence of CPP-like motifs within specific lineages.

**Figure 1.**
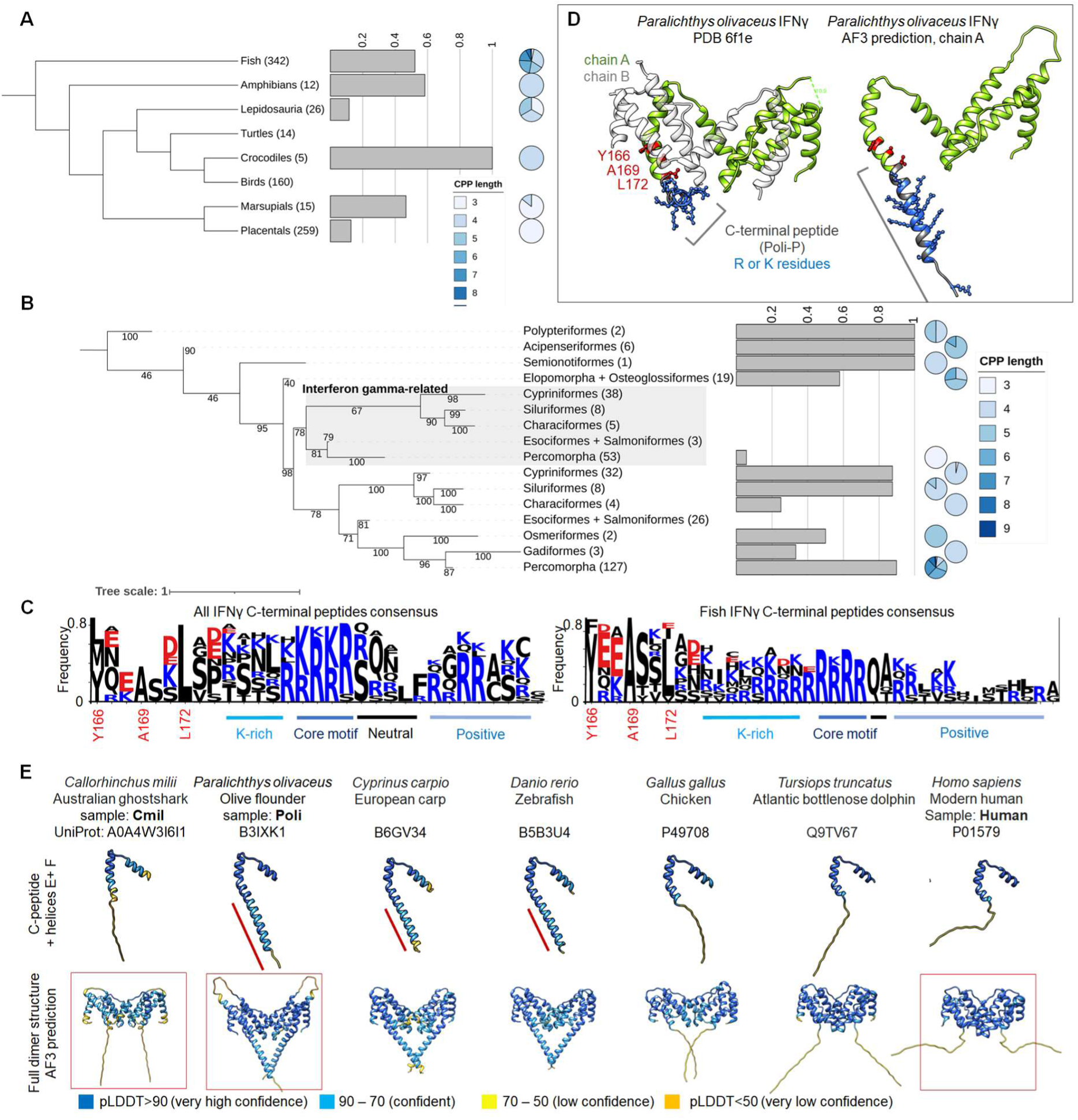
**-Presence, diversity, occurrence, and structure of arginine-rich CPP-like motifs among IFN-γ proteins**. (A) Cladogram illustrating vertebrate evolutionary relationships. To the right of each group, a bar plot shows the percentage of IFNγ sequences containing a CPP-like motif, and the adjacent pie chart depicts the length of these motifs. The number in parentheses indicates the total number of IFNγ sequences identified in each group. (B) Maximum-likelihood phylogenetic tree (supported by 1000 bootstrap replicates) showing the evolution of IFNγ sequences in fish. The numbers in parentheses indicate the total number of IFNγ sequences identified in each order. The bar plots and pie charts depict the prevalence and length of the CPP-like motifs, as in part (A). (C) Sequence logos of the C-terminal region of canonical IFN-γ proteins generated from multiple sequence alignments, for all chordates (left) and fish (right). Conserved residues at the end of helix 6 (Y166, A169, and L172; numbering according to *Paralichthys olivaceus* IFN-γ, PDB 6f1e) were used as alignment anchors. Amino acids occurring at frequencies below 5% were omitted. (D) Structural organisation of the C-terminal region of *Paralichthys olivaceus* IFN-γ. Left, experimental crystal structure (PDB 6f1e) showing the initiation of an additional C-terminal α-helix (in blue) in one subunit of the homodimer (chain A). Right, AlphaFold3 prediction of the same protein showing a continuous α-helical extension of helix 6 encompassing the entire C-terminal peptide. (E) AlphaFold3 predicted structures of representative IFN-γ proteins from diverse vertebrate lineages illustrating two distinct classes of C-terminal organisation. Representative proteins selected for subsequent experimental analyses are indicated by red frames.

Our analysis revealed marked lineage-specific differences in the prevalence and length of C-terminal CPP-like motifs, most of which consisted of consecutive stretches of arginine residues. Bony fish exhibited the greatest diversity and the longest CPP-like motifs. In contrast, both motif frequency and length generally decreased across tetrapod lineages (Fig. 1A). Among placental mammals, CPP-like motifs were restricted to a few groups, namely hystricomorph rodents (e.g. blesmols and guinea pigs), new world leaf-nosed bats (*Phyllostomidae*), and cetaceans (Supp. Fig. S1). The presence of these motifs in aquatic tetrapods suggests that retention of the C-terminal region may be associated with adaptation to aquatic environments.

In contrast, all fish IFN-γrel proteins consistently lacked the characteristic C-terminal peptide (Fig. 1B), further supporting their functional divergence from canonical IFN-γ.

For further analyses of IFNγ C-terminal peptides, we used the previously identified structure of IFNγ from *Paralichthys olivaceus* (PDB 6f1e) and the almost absolutely conserved residues identified in our multiple sequence alignments at the end of helix 6: tyrosine 166, alanine 169, and leucine 172 (numbering according to PDB 6f1e) to create representative sequence logos in Fig. 1C, D. The previously reported conserved consensus motif KRKR [21] was re-evaluated using our expanded dataset, revealing that it is less strictly conserved than originally suggested. Furthermore, the pattern appears more complex: the R/K-R/K-R/K-R sequence represents only the core of a broader motif. This extended motif typically begins with a lysine-rich stretch, followed by an increasing number of arginine residues, then the core sequence itself, and concludes with 2–5 neutral residues ending in a peptide region enriched in positively charged amino acids (Fig. 1C, 1D).

### Structural modelling reveals distinct conformational states of IFN-γ C-termini

To investigate whether IFN-γ C-termini sequence differences among chordate groups are accompanied by structural variation, we first examined the available experimental structures. Among all currently available IFN-γ structures, the crystal structure of *P. olivaceus* IFN-γ is unique in displaying the beginning of an additional α-helix formed by the C-terminal peptide in one subunit of the homodimer (chain A, Fig. 1D). Such an arrangement has not been observed in any other experimentally determined IFN-γ structure. Consistent with this observation, AlphaFold3 predicted the entire C-terminal region of *P. olivaceus* IFN-γ to adopt a continuous α-helical conformation extending helix 6 (Fig. 1D). In contrast, the corresponding prediction for human IFN-γ indicated that the C-terminal peptide remains intrinsically disordered (Fig. 1E).

We next explored whether these structural differences represent isolated cases or reflect broader evolutionary patterns. We therefore generated AlphaFold3 models [34] for representative IFN-γ proteins spanning the major vertebrate lineages (Fig. 1E). The predicted structures could be broadly classified into two groups. The first group, represented by *P. olivaceus*, common carp (*Cyprinus carpio*), and zebrafish (*Danio rerio*), contained a structured C-terminal extension that formed a characteristic “heart-shaped” arrangement together with helix 6 (Fig. 1E). The second group comprised proteins with an unstructured C-terminus, including human, chicken, and the cartilaginous fish Australian ghostshark (*Callorhinchus milii*). The presence of both structural classes in evolutionarily distant vertebrate lineages indicates that C-terminal organisation does not simply follow phylogenetic relationships.

Based on these analyses, we selected representative IFN-γ proteins for subsequent experimental characterisation. *P. olivaceus* IFN-γ (PoliIFN-γ) was chosen as a representative of the structured C-terminal class. In contrast, Australian ghostshark IFN-γ (CmilIFN-γ) and human IFN-γ (hIFN-γ) were selected as representatives of the unstructured class. Their isolated C-terminal peptides, hereafter referred to as Poli, Cmil, and Human, respectively, were used in subsequent computational, biophysical, and cellular experiments. Nonaarginine (R_9_) and tetraarginine (R_4_) serve as positive and negative controls for cell internalisation, respectively. Additionally, nonalysine (K_9_) was used as an example of a peptide that does not interact with the POPC lipid membrane.

### Molecular dynamics suggests membrane interaction of IFN-γ C-terminal peptides

The evolutionary conservation of highly positively charged C-terminal sequences together with their predicted structural diversity prompted us to investigate whether these peptides differ in their intrinsic ability to interact with lipid membranes. To examine the molecular basis of these interactions, we performed all-atom molecular dynamics (MD) simulations of representative Poli, Cmil, Human, and R_9_ peptides in the presence of two model phospholipid bilayers: neutral zwitterionic POPC (phosphatidylcholine) and a negatively charged POPC/DOPS (dioleoylphosphatidylcholine, dioleoylphosphatidylserine) 80/20 mixture (details are given in the Materials and Methods section). Simulations were carried out using the GROMACS program package and two independent force fields, the standard CHARMM36/LJ-PME and the charge-scaled ProsECCo force field [35], to ensure that the observed interactions were robust and independent of a particular force field.

The results of the MD simulations are depicted in Fig. 2. Figure 2A shows a representative snapshot of the simulation system, consisting of an upright-oriented lipid membrane with two peptides in close contact with the lipid bilayer, surrounded by a water and ion environment. In addition, the panel shows an example density profile calculated along the axis perpendicular to the membrane. Fig. 2B provides examples of peptides associating with the POPC membrane (R_9_, Poli) and a peptide that stays in the bulk solution without interacting with the membrane (K_9_). Figure 2C shows the density profiles calculated from two distinct simulation setups: one containing a neutral POPC membrane (top) and the other containing a negatively charged POPC/DOPS 80/20 membrane (bottom).

**Figure 2.**
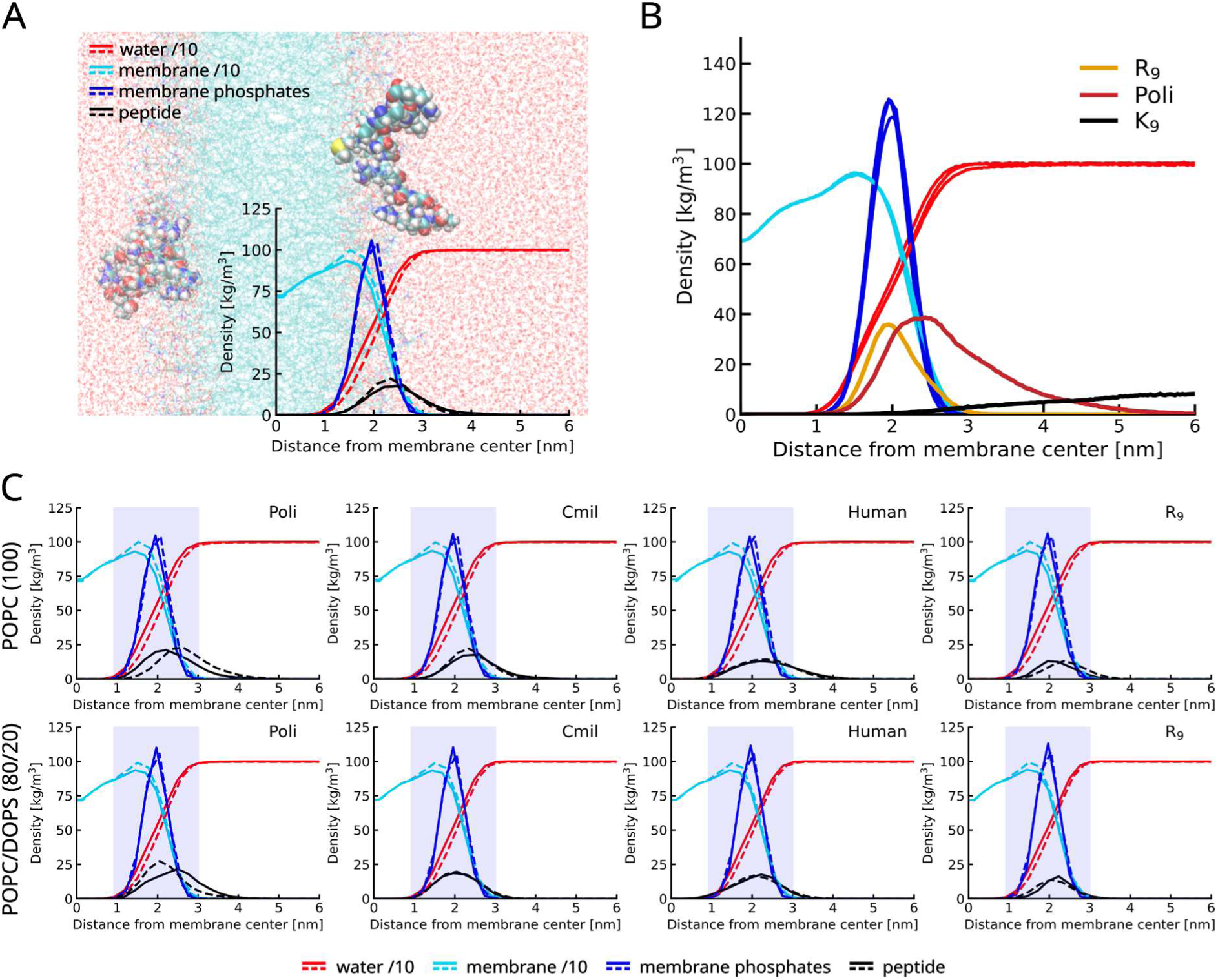
-Molecular dynamics simulations of IFN-γ C-terminal peptides interacting with lipid membranes. (A) Illustrative image of simulation setup. Water molecules are rendered in red, a perpendicularly oriented membrane in cyan, and two mixed-colour peptides (Cmil) interact with the membrane. In the bottom right corner, a density graph shows the densities of specific groups as a function of their distance from the membrane centre plane, in accordance with the underlying illustration. (B) Example of peptides interacting with the POPC membrane (R_9_, Poli), and a peptide not adhering to the membrane surface (K_9_), instead staying in the aqueous bulk. Data are from another simulation replica set. (C) Specific group densities as a function of distance from the membrane centre for both the POPC membrane (top part) and POPC/DOPS 80/20 membrane (bottom part), for the Poli, Cmil, Human, and R_9_ peptides (from right to left). Solid lines represent MD simulation results using the CHARMM force field, and dashed lines using the charge-scaled ProsECCo force field. The light blue region represents the membrane boundary (its phosphate region). For clarity in the image, we scaled the water and membrane densities by a factor of 10 (denoted as “water /10” and “membrane/10” in the legend).

Despite differences in sequence and structure, all tested IFN-γ C-terminal peptides interacted with lipid bilayers, regardless of membrane charge or force field used. Their density profiles consistently showed peaks near the membrane phosphate region, indicating association with the membrane (Fig. 2C).

### IFN-γ C-terminal peptides interact preferentially with negatively charged lipid membranes

To experimentally assess the membrane-interacting properties of the studied peptides, we prepared Large unilamellar vesicles (LUVs) with a defined composition of either net-neutral-charge POPC or a negatively charged DOPC/DOPS 80/20 lipid mixture. We monitored peptide interactions with these artificial lipid membranes using Laurdan fluorescence spectroscopy and Dynamic Light Scattering (DLS).

The emission spectrum of the fluorescent dye Laurdan is sensitive to the polarity and viscosity of its environment. Laurdan, located at the carbonyl level of the phospholipid bilayer, senses the hydrogen bond network formed by water molecules bound to the lipids at this level, thereby reporting local lipid packing and hydration [36–38]. Importantly, Laurdan fluorescence is sensitive to membrane changes induced by peripheral and integral peptide binding, as demonstrated previously [39, 40]. To observe the kinetics of the peptide binding to the membrane, we used the Laurdan generalized polarization (GP) measured after 1, 10, 20 and 30 minutes after peptide addition to LUVs (Fig. 3A, B) No changes in GP were observed for POPC liposomes for any of the peptides, indicating none or weak peptide-membrane interaction (Fig. 3A). On the other hand, the increase in GP values in time suggests Poli and R_9_ peptides bind to the DOPC/DOPS membrane (Fig. 3B). While still exhibiting a non-negligible GP change, the Cmil and Human peptide-membrane interaction is weaker than that of Poli and R_9_.

**Figure 3.**
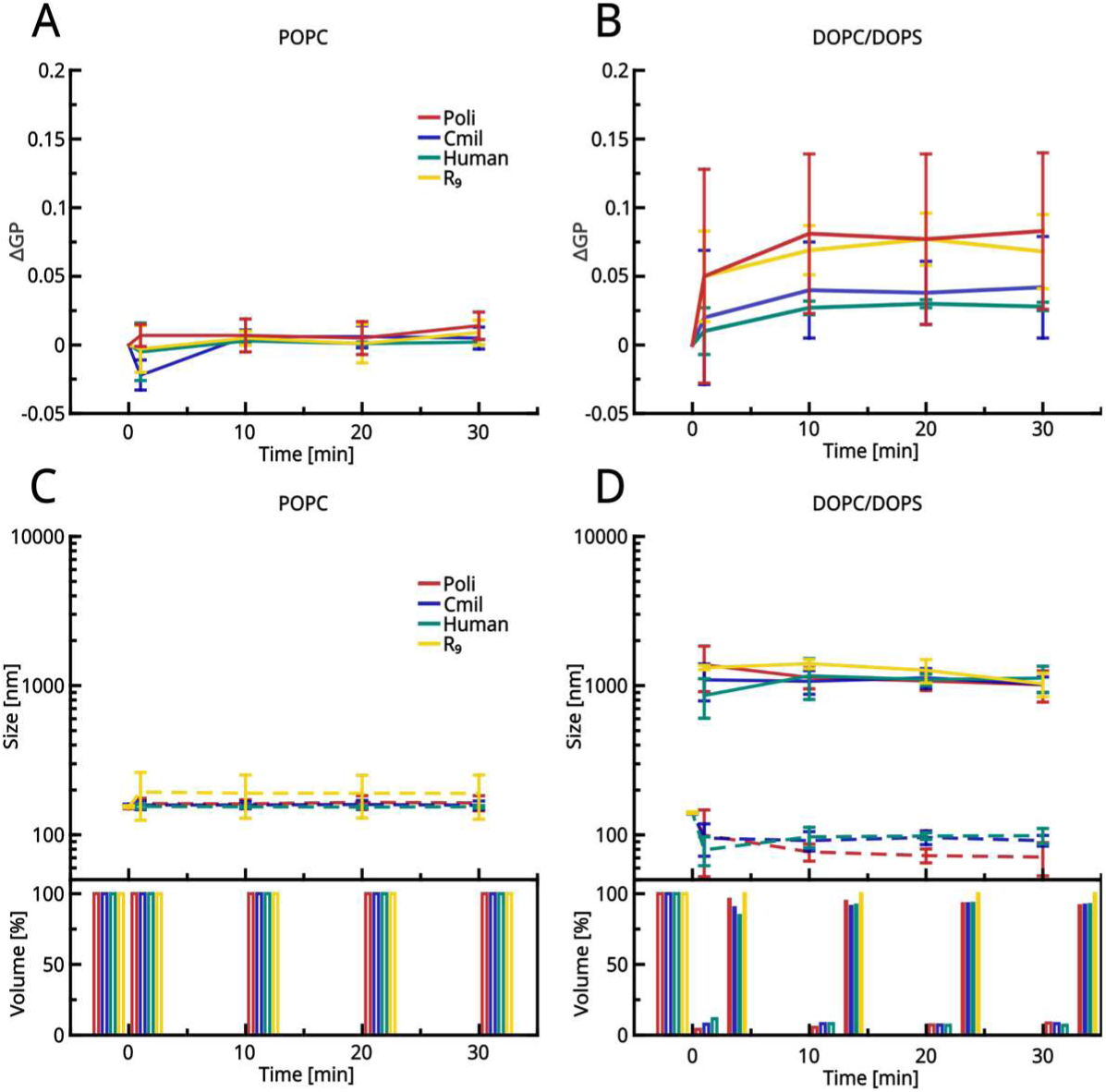
-IFN-γ C-terminal peptides interact with LUV lipid membrane. Top: Peptide-lipid interaction measured as a change in Laurdan GP after peptide addition to the POPC LUVs (A) or DOPC/DOPS 80/20 mixture LUVs (B). The initial value, before peptide addition, is denoted as time 0 min. Time point 1 min represents the measurement right after peptide addition. Bottom: Vesicle size measurements by DLS: POPC (C) and DOPC/DOPS LUVs (D). Similarly, time point 0 min denotes the measurement immediately before peptide addition, and 1 min denotes the measurement immediately after. The “vesicles” population (up to 300 nm) is represented by dashed lines, while full lines denote the “aggregates” population (∼1 μm). Bar plots at the bottom show percentage representations of observed DLS size populations (empty bars: small population, full bars: big population). All measurements were carried out in triplicate.

DLS was used to assess more pronounced changes that perturb membrane organisation and induce vesicle aggregation, as evidenced by changes in vesicle size and distribution. Average LUV size after peptide addition was monitored over the same time course as the GP measurements (Fig. 3C, D). Consistently with Laurdan GP, DLS revealed no substantial change in the size of POPC-based LUVs following the peptide addition (Fig. 3C). In contrast, a rapid vesicle change is visible right after peptide addition to the DOPC/DOPS vesicle suspension (Fig. 3D). Here, two distinct sample populations arise: the “vesicles” population with a size of up to 300 nm, which most likely corresponds to unaffected LUV vesicles (or possibly small parts of burst vesicles), and the “aggregates” population, that represents large vesicle aggregates (around 1 μm in size). This population arises for all examined peptides with DOPC/DOPS LUVs, indicating that these peptides induce notable LUV aggregation, with very few vesicles unaffected (Fig. 3D).

Overall, the DLS and Laurdan spectroscopy experiments show little to no interaction between the investigated peptides and the POPC LUVs In contrast, all investigated peptides interact with the negatively charged DOPC/DOPS 80/20 lipid vesicles. Poli and R_9_ peptides showed significantly stronger signals than Cmil or Human peptides. These results are in contrast to the MD predictions that all peptides interact with both neutral and negatively charged membranes. We assume that peptide interactions with the POPC membrane may be too weak to be detected by Laurdan. Additionally, as a consequence of weak peptide-membrane interactions, DLS might not report LUV aggregation or any other vesicle size change.

### IFN-γ C-terminal peptides interact with living cells

To further examine the interactions of IFN-γ orthologue C-terminal peptides with biological membranes, we followed their association with U-2 OS cells using live-cell confocal microscopy and flow cytometry.

Peptides labelled with the fluorescent dye Tamra were incubated with U-2 OS cells at a final concentration of 10 μM and monitored for 60 min. Tamra dye was included as a control for free dye internalisation. Confocal images of cells incubated with peptides for 10 mins are shown in Fig. 4A. We observed a strong increase in fluorescence intensity within cells treated with R_9_ and Poli peptides. Poli rapidly penetrated all cells (Fig. 4) and induced cell death in a subset of the population within 10 min of incubation. The rapid progression of this cytotoxic effect is further illustrated by time-series images acquired after peptide addition, with most cells showing loss of viability by 15 min (Fig. S2A). Due to the similar patterns observed for Poli and R_9_ cell internalisation, we propose that Poli might enter cells via a passive mechanism, without the need of ATP, similarly to R_9_. Consistent with this possibility, a similar internalisation pattern was observed when Poli was fused to the C-terminus of the fluorescent protein UnaG, although the process was considerably slower than that observed with the Tamra-labelled peptide (Fig. S2B). Human peptides associated with the cell membrane, forming punctate fluorescence signals, and also penetrated into the cell interior, although to a much lesser extent than the Poli or R_9_ peptides. Notably, the formation of similar puncta, corresponding to peptide-induced membrane stacking, has been identified as a key step in the passive penetration of peptides into cells [40]. In contrast, Cmil showed only limited internalisation and remained predominantly outside the cells.

**Figure 4.**
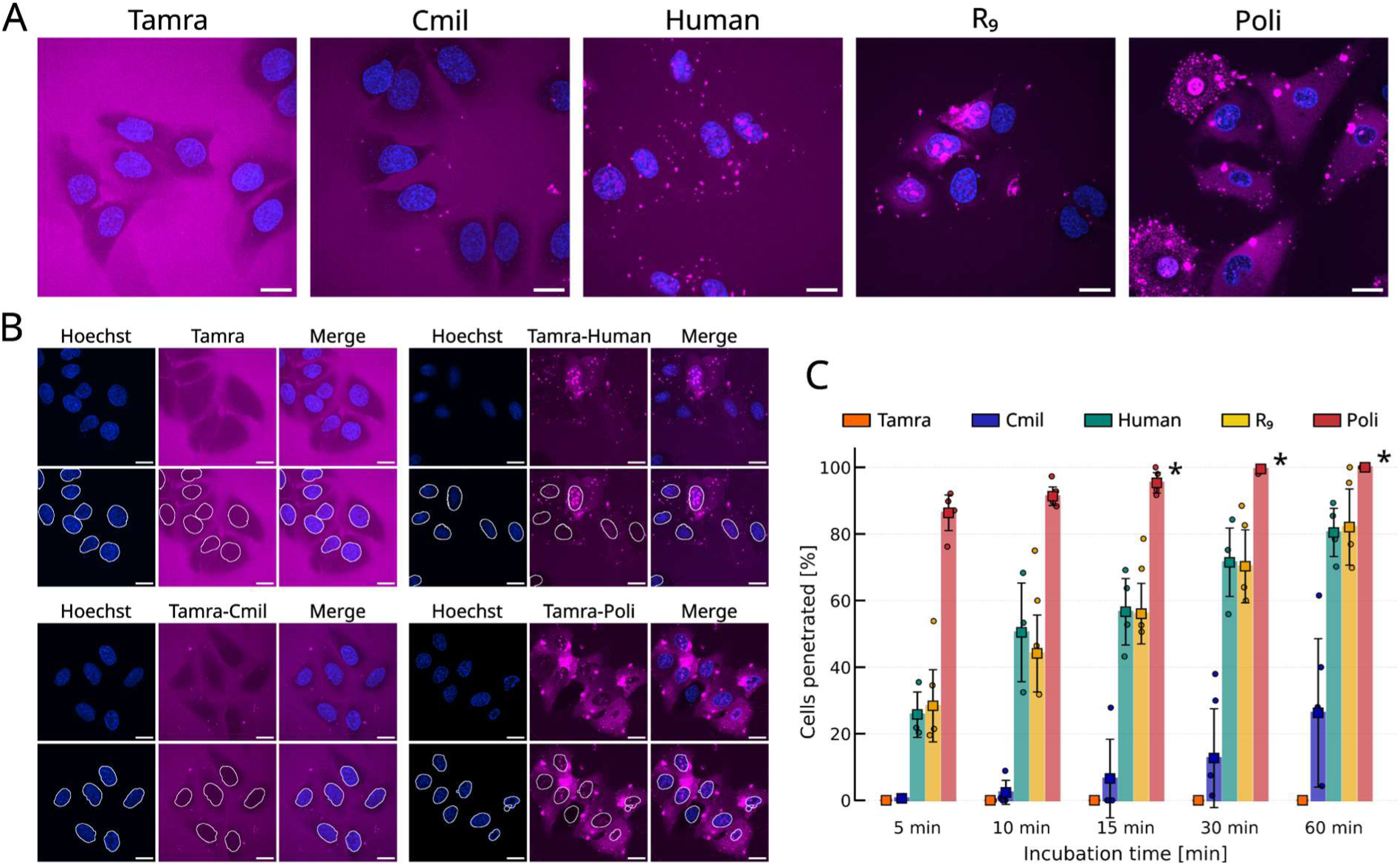
-Tamra-labelled IFN-γ orthologue C-terminal peptides penetrate U-2 OS cells. (A) Illustrative confocal live-cell imaging snapshots of 10 μM Tamra-labelled peptides (red) at 10 min cell incubation time. Cell nuclei are labelled with Hoechst (blue). The scale bar is 20 μm. (B) Since peptides localise within the cell nucleus shortly after entering the cell, to determine whether the cell is penetrated, the average Tamra-channel fluorescence intensity within the cell nuclei was compared with the average outer-space fluorescence intensity. To illustrate this workflow, four panels (Tamra dye, Human, Cmil, and Poli) show snapshots from a 5 min peptide-cell incubation. Each panel shows the Hoechst and Tamra channels and the Merge (top part), with the cell nuclei region borders emphasised for these channels (bottom). The scale bar is 20 μm. (C) Weighted average of the percentage of penetrated cells during a 60 min incubation, calculated from at least 3 independent replicates. The asterisks (*) at the Poli incubation of 15 min, 30 min, and 60 min denote the observation that most of the cells here are peptide-penetrated and dead.

Furthermore, shortly after peptides enter the cell interior, we observed their nuclear localisation. In the case of Poli, R_9_, and Human, strong nuclear accumulation is detectable already after 5 min (Fig. 4B). After 60 min, all studied peptides were detected in the nucleus, with nuclear penetration observed in 100% of cells for Poli, more than 80% of cells for R_9_ and the Human peptide, and approximately 30% for Cmil (Fig. 4C).

To obtain a quantitative assessment of peptide-cell interactions, we performed flow cytometry measurements using Tamra-labelled peptides across a range of several concentrations. Tamra-labelled R_4_ and R_9_ peptides were included as controls representing weakly interacting and strongly interacting CPPs, respectively [40]. Individual cells were selected using a two-step gating strategy (Supplementary Fig. S3). Peptides exhibiting membrane association or cellular uptake produced concentration-dependent rightward shifts in fluorescence intensity distributions, accompanied by reduced relative background contribution, providing a robust measure of peptide-cell interaction strength.

All studied IFN-γ C-terminal peptides exhibited measurable interactions with U-2 OS cells, although with markedly different efficiencies (Fig. 5A). As expected, the positive control peptide R_9_ induced a pronounced fluorescence shift, whereas the negative control R_4_ produced only a minimal shift (Fig. 5B), reaching approximately 24% of the mean PE-A fluorescence measured for R_9_ at a concentration of 15 µM (Fig. 5C). Among the IFN-γ C-terminal peptides, Cmil exhibited the weakest cellular association, reaching approximately 53% of the R_9_ fluorescence intensity, while the Human peptide reached approximately 79%. In contrast, the Poli produced exceptionally strong fluorescence signals, with a mean intensity nearly 10-fold higher than that of R_9_. These findings are consistent with the confocal microscopy experiments. The mean and median fluorescence intensities are provided in Table S1.

**Figure 5.**
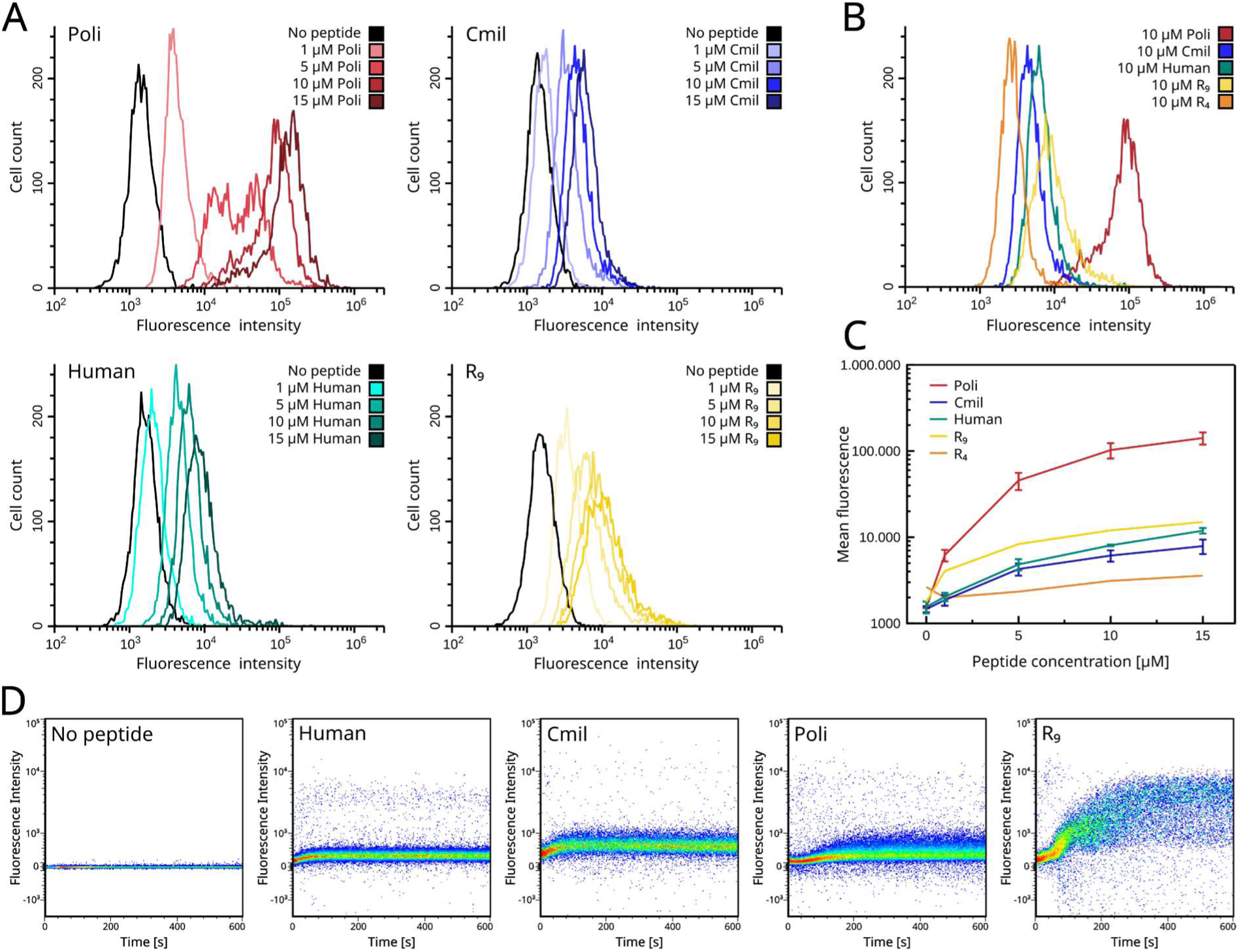
-Association of IFN-γ orthologue C-terminal Tamra-labelled peptides with human osteosarcoma U-2 OS cells. Human U-2 OS cells were incubated with Tamra-labelled peptides for 20 min and analysed by flow cytometry. (A) Representative PE-A fluorescence intensity histograms following incubation with peptides at 1, 5, 10, and 15 μM. (B) Overlay of fluorescence intensity histograms comparing IFN-γ orthologue C-terminal peptides with the control peptides R_4_ and R_9_ at 10 μM. (C) Mean fluorescence intensity of cells incubated with the indicated peptide concentrations. Data for IFN-γ orthologue C-terminal peptides represent three independent biological replicates; R_4_ and R_9_ control peptides were measured once. Error bars indicate SD. (D) Time-resolved flow cytometry analysis after adding 1 μM peptide at time 0. At this concentration, all IFN-γ orthologue C-terminal peptides exhibited comparable levels of cellular association. The R_4_ and R_9_ control dataset is reproduced from Baxova et al. (2026) [40] with permission and included for direct comparison, whereas all other data are newly generated in this study.

To observe the evolution of cell-associated Tamra-labelled fluorescence over time, we performed time-dependent flow cytometry experiments using 1 µM peptides. We measured fluorescence intensity immediately after adding the peptide to the U-2 OS cell suspension. The gating strategy is shown in Supporting Fig. S4. Here, cell events were divided into 3 populations: singlets (the majority), aggregates, and subcellular fragments. Fig. 5D shows detected fluorescence of singlets. Generally, increases in cell-event fluorescence are comparable for Poli, Cmil, and Human when incubated with 1 µM peptide. In contrast, R9-associated fluorescence increased markedly even at this peptide concentration, whereas the Poli peptide did not exhibit similarly strong cell association.

Additionally, time-resolved flow cytometry measurements in the absence of cells confirmed that the peptides themselves did not form large aggregates (see Fig. S5).

Interestingly, additional observations from exploratory AlphaFold 3 predictions raise the possibility that the human IFNγ C-terminus exhibits lipid-dependent conformational plasticity (Fig. 6). While the C-terminal region was predicted to be largely disordered in the absence of lipids, inclusion of either palmitic acid or DOPS resulted in the prediction of an α-helical conformation positioned in association with the lipid molecules, albeit with low or very low pLDDT scores (3 replicates, 5 independent seeds). The convergence of the fatty acid and lipid-containing predictions is intriguing. Such a conformational transition would be consistent with the behaviour of secondary amphipathic cell-penetrating peptides, some of which can undergo a disorder-to-helix transition upon interacting with phospholipid membranes [41, 42]. In this context, previous studies have demonstrated the preferential association of whole hIFNγ protein with negatively charged liposomes [25], although no substantial changes in the overall protein conformation were detected upon lipid association [43]. While the limited confidence of the AlphaFold3 predictions and the absence of an explicit membrane environment preclude definitive structural interpretation [34], these observations raise the possibility that the IFNγ C-terminus possesses an intrinsic lipid-responsive helical propensity that may contribute to its interactions with cellular membranes and experimental observations. This possibility is further supported by the presence of a predicted helical conformation in the C-terminal regions of IFNγ orthologs (Fig. 1E), as well as by the experimentally determined structure of human IFNγ (PDB 1eku).

**Figure 6.**
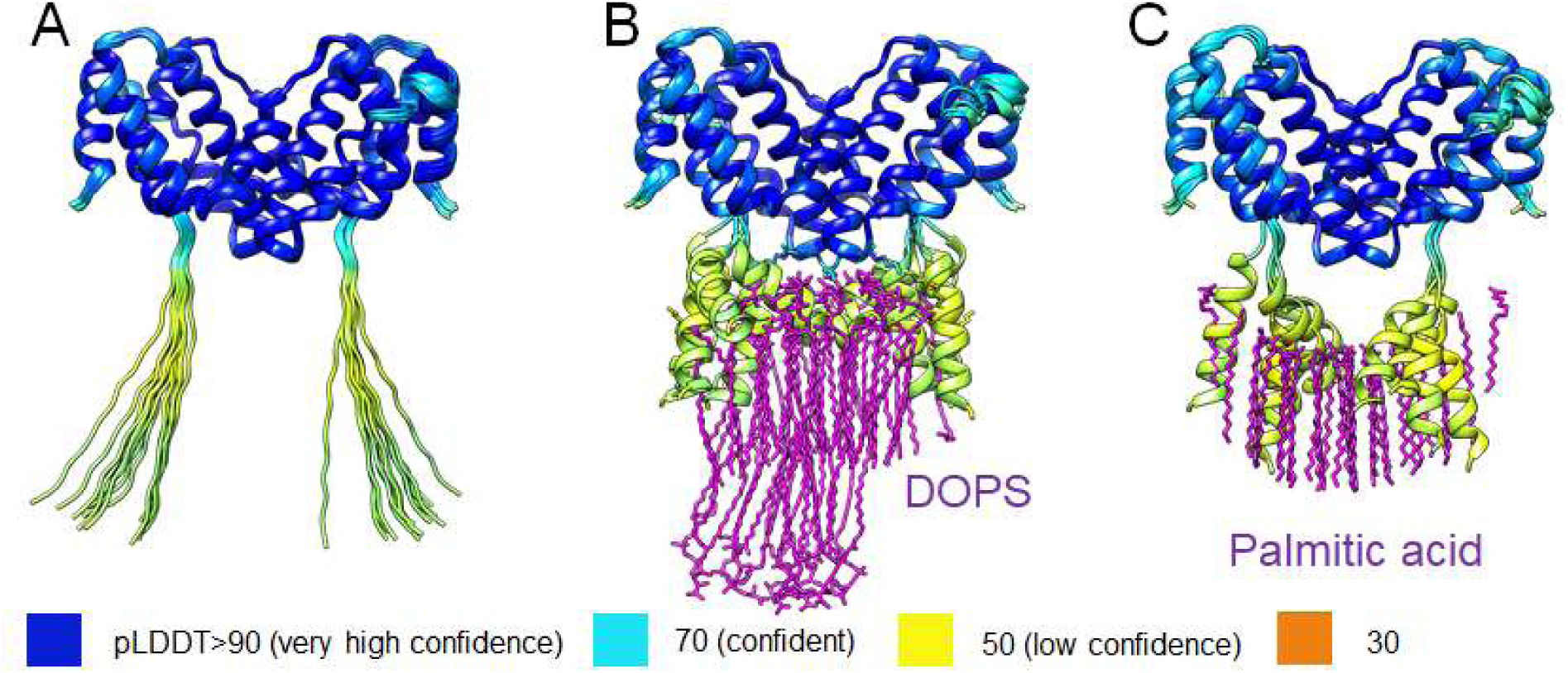
Lipid/fatty-acid-associated α-helical conformation of the human IFNγ C-terminus predicted by AlphaFold3. (A) In the absence of lipids, the C-terminal region is predicted to adopt a largely disordered conformation. (B, C) Inclusion of 30 molecules of dioleoylphosphatidylserine (DOPS) or palmitic acid, respectively, results in the prediction of an α-helical C-terminal conformation associated with the lipid molecules. Structures are coloured using a continuous scale based on the predicted local distance difference test (pLDDT). The predicted conformations have relatively low confidence and should therefore be interpreted as exploratory behaviour suggesting models rather than evidence of a defined lipid-bound structure.

## Conclusions

IFN-γ is one of the most extensively studied immune cytokines, with established roles in immune regulation, antimicrobial defence, and antitumour immunity. Beyond its canonical signalling functions, its positively charged C-terminal region has been associated with nuclear localisation and heparan sulfate binding. Here, we investigated the membrane-interacting properties of this region by combining comparative sequence and phylogeny analyses, molecular dynamics simulations, biophysical measurements, live-cell microscopy, and FACS interaction assays.

Analysis of 833 IFN-γ sequences revealed that the C-terminal motif is more complex and evolutionarily variable than the previously described KRKR consensus. Rather than a universally conserved short sequence, this region exhibits a broader pattern of positively charged residues, characterised by an initial lysine-rich stretch, an increasing contribution of arginine residues, a central basic motif with R/K-R/K-R/K-R sequence, and a variable C-terminal segment containing neutral residues within a positively charged context. The new motif captures sequence and biochemical peptide diversity beyond tetrapods and is applicable to all IFN-γ sequences.

To isolate the intrinsic properties of this region from the structural and functional contributions of the rest of the biophysically difficult-to-study cytokine, we deliberately focused on chemically synthesised, sequence-authenticated C-terminal peptides representing three distinct vertebrate orthologues: human IFN-γ, the arginine-rich *P. olivaceus*, and phylogenetically basal *C. milii* orthologues.

Molecular dynamics simulations indicated that all three C-terminal peptides can interact with both zwitterionic POPC and negatively charged lipid membranes. Consistent with these predictions, Laurdan fluorescence spectroscopy and DLS experiments with LUVs demonstrated interactions with negatively charged DOPC/DOPS membranes. On the other hand, despite the simulation-predicted interactions with POPC, we observe no Laurdan GP change and no peptide-induced LUV aggregation, suggesting a weak or no peptide-POPC membrane interaction for all the investigated peptides. Live-cell imaging further revealed distinct cellular interaction patterns among the peptides, with the arginine-rich Poli peptide exhibiting the strongest observed interactions and possible passive cell penetration, exceeding those of the canonical cell-penetrating peptide R_9_ under our experimental conditions. These findings predict substantial functional differences among evolutionarily related IFN-γ C-terminal regions and suggest that their amino acid composition modulates their interactions with both model membranes and living cells.

Together, our results identify the IFN-γ C-terminus as an evolutionarily diverse, positively charged region with intrinsic membrane-interacting properties. In at least some orthologues, these properties may facilitate concentration-dependent interactions with, and potentially translocation across cellular membranes.

Although the precise mechanism of translocation remains to be elucidated, these observations suggest that the C-terminus contributes to cytokine distribution, local retention, or engagement with cell-surface components in addition to its previously described functions. Given the importance of IFN-γ oligomerisation and receptor-complex stoichiometry for signalling, membrane association could also influence the local availability and spatial organisation of the cytokine. Furthermore, since IFN-γ is a direct target of viral immunomodulatory proteins, the ability of its C-terminal region to promote membrane translocation raises the intriguing possibility that this mechanism could serve as a direct countermeasure, allowing signalling even in the presence of such proteins. Our findings thus reveal an important, previously unrecognised aspect of IFN-γ biology and provide a robust foundation for future investigations of how the membrane-interacting properties of its evolutionarily diverse C-termini influence cytokine function in immune and host-pathogen contexts.

## Materials and Methods

### Sequence retrieval, filtering, and alignment

We queried the NCBI Protein database for interferon gamma protein sequences in chordates using the following search string: ("interferon gamma"[Protein Name] OR "interferon gamma related"[Protein Name]) AND "chordates"[porgn: txid7711] NOT "partial" NOT "receptor" NOT "precursor" NOT "low quality" NOT "inducible" NOT "hypothetical". This returned 824 protein sequences: 129 from fish and 695 from other chordates. To expand the dataset, we performed a BLASTP search against the NCBI non-redundant (nr) protein database, restricted to the Chordata taxon (taxid:7711). Only hits that covered at least 80% of the query sequence were retained. For fish sequences, this process was iterated for four rounds (the final round yielded no additional sequences). For other chordates, only one iteration was performed due to the increasing presence of false positives in subsequent rounds. The final dataset comprised 464 bony fish IFNγ sequences and 1076 IFNγ sequences from other chordates. All retrieved sequences were aligned using MUSCLE with default parameters for protein sequences. To reduce redundancy, we removed sequences from the same species that differed in at most 3% of aligned positions, excluding sites where both sequences had gaps. Additional manual curation was performed to eliminate incomplete or likely misannotated sequences. The resulting dataset consisted of 342 bony fish and 491 non-fish chordate IFNγ sequences.

### CPP detection

To identify CPP motifs within IFNγ proteins, we downloaded all known L-chirality CPP sequences from the CPPsite 2.0 database (http://crdd.osdd.net/raghava/cppsite/), totalling 1564 unique sequences. Each IFNγ sequence was scanned for exact matches to these CPP motifs within the C-terminal 20 amino acids. If multiple CPPs matched, the longest match was retained.

### Phylogenetic analysis

Phylogenetic trees were constructed for four chordate groups: bony fish (342 sequences), rodents (46), bats (25), and artiodactyls (58). Protein sequences were aligned using Clustal Omega (v1.2.4) with default settings. Alignments were trimmed using trimAl (v1.5.rev0) with the -automated1 option. Maximum-likelihood trees were inferred using IQ-TREE (v2.3.6) with automatic model selection and 1000 ultrafast bootstrap replicates (-m TEST -B 1000). For the rodent, bat, and artiodactyl trees, the cat (*Felis catus*) IFNγ sequence was included as an outgroup to root the trees. Trees were visualised with iTOL, and branches with bootstrap support below 50% were collapsed.

Sequence logos were generated using the Python Logomaker package from the C-terminal regions of the aligned IFNγ sequences to visualise residue conservation and variability. Amino acids were individually colored according to the ‘dmslogo_chargé colour scheme, with histidine specifically recoloured in black. Residues with a frequency below 5% were excluded from visualisation. Particular attention was given to the positions corresponding to Y166, A169, and L172 (numbering according to PDB 6F1E), which are highly conserved across the analysed sequences.

### MD simulation details

Molecular dynamics simulations were performed using the Gromacs2022 program package. Simulation systems were built with CHARMM-GUI [44]. They are composed of a lipid membrane containing 256 lipids per leaflet, 100 waters per lipid, and 2 peptide molecules. 150 mM KCl was added to mimic a biological system, with additional potassium or chloride ions to maintain electroneutrality. Two Poli, Human, Cmil, or R_9_ peptides were simulated using either a neutral 100% POPC membrane or a lipid composition of 80% POPC and 20% DOPS, negatively charged. Initially, peptides were positioned sufficiently far from the membrane within the simulation box. Two force-field parameter sets were used: CHARMM36/LJ-PME and a charge-scaled ProsECCo [35].

The isobaric-isothermal (NpT) ensemble was used. Nose-Hoover thermostat [45] was set at 303.15 ℃ with a 1 ps time constant for the groups of membrane, proteins, and water with ions. Semiisotropic Parrinello-Rahman pressure coupling [46] was set at 1 bar. Long-range interactions were treated with Particle-Mesh Ewald electrostatics (PME, [47]) with a 1.2 nm cutoff. Water molecules were described by the TIP3P water model [48] with SETTLE constraints [49]. Other hydrogen atom bonds were constrained by the LINCS algorithm [50]. 1 µs simulation runs were preequilibrated for 10 ns, and a 2 fs leap-frog integration timestep was used. The resulting symmetrised density profiles were calculated from the last 200 ns of the simulation runs.

### Reagents

POPC (1-palmitoyl-2-oleoyl-glycero-3-phosphocholine), DOPC (1,2-dioleoyl-sn-glycero-3-phosphocholine), and DOPS (1,2-dioleoyl-sn-glycero-3-phospho-L-serine) lipids dissolved in chloroform (Avanti Polar Lipids, USA); Laurdan dissolved in methanol (min 99,8 %, p.a., Carl Roth, Germany) Peptides Poli (NTKMQRRRRRRRRQARKVKTPTRA) with a positive charge of +13, Cmil (NVNGRRRRRSNMGGRRRSRT) with a charge of +9, Human (AKTGKRKRSQMLFRGRRASQ) charged +8, and R_9_ (RRRRRRRRR) with a charge of +9, as well as their Tamra-labelled counterparts, were synthesised in-house at IOCB, Prague.

### LUV preparation

Lipids dissolved in chloroform were prepared at a target concentration of 1 mM with 1% Laurdan added, evaporated under argon flux, and left under vacuum overnight. The lipid film was rehydrated with PBS +/+, briefly vortexed, and then extruded 21 times through a 100 nm pore-size membrane filter (Whatman). For both lipid compositions used, neutral POPC (100%) and negatively charged DOPC/DOPS (80%/20%), three separate sample replicates were prepared and measured, adding peptides at a peptide/lipid ratio of 1/1.

### DLS measurements

Samples with a total volume of 200 μl were measured in a UV-Cuvette micro (Brand) using a Malvern Zetasizer Nano ZS at 25℃. The LUV size was measured immediately before peptide addition (designated as time 0 min), immediately after peptide addition (designated as time 1 min), and subsequently at 10 min, 20 min, and 30 min after peptide addition, all in 3 technical replicates. If one of these replicates included a size peak not observed in the other two, it was considered an impurity and excluded from data processing. Samples were stirred before each measurement.

### Laurdan spectra measurements

In order to obtain the General polarisation factor (GP), defined by the spectral intensities at 440 nm (I_440_) and 490 nm (I_490_) as: GP=(I_440_-I_490_)/(I_440_+I_490_), Laurdan emission spectra were measured (PicoQuant Fluotime 300). Excited at 374 nm, the spectra were obtained in the range of 400-580 nm. Measurements were performed at 24℃ after a minimum of 10 minutes of thermal equilibration. Due to the possibility of aggregate formation (as indicated by the DLS measurement results), samples were thoroughly stirred before each measurement.

For each sample, spectra were measured immediately before peptide addition (designated as time 0 min), immediately after peptide addition (designated as time 1 min), and subsequently at 10 min, 20 min, and 30 min after peptide addition, in 3 technical replicates.

### Cell culture

Human bone osteosarcoma (U-2 OS) cells were cultured at 37℃ in a 5% CO_2_ atmosphere in high-glucose DMEM media (Sigma-Aldrich) supplemented with 10% FBS and 1% penicillin-streptomycin.

### Live cell imaging

U-2 OS cells were transferred into Ibidi μ-slide 8-well glass-bottom plates (approximately 45,000 cells/well) and incubated overnight. After washing with PBS, cells were incubated with 1:1000 Hoechst in PBS staining solution for 5-10 min, washed 3x with PBS, and further incubated in 250 µl Hank’s Balanced Salt Solution buffer (HBSS +/+, Gibco).

Nikon CSU-W1 (based on an inverted widefield microscope Nikon Eclipse Ti2-E), equipped with an OKOLab Microscope Incubator and a 60x NA 1.20 water-immersion objective (CF Plan Apo VC, Nikon), was used for live-cell imaging. Cells were incubated at 37℃ in a 5% CO_2_ atmosphere during the measurements.

We captured cell nuclei with Hoechst dye using a confocal 405 nm laser channel with 50-200 ms exposure time and 1.4-4.8 mW laser power. We imaged Tamra-labelled peptides using the 561 nm confocal channel with an exposure of 50-100 ms and a laser power of 0.6-1.2 mW. All images shown have been adjusted for brightness and contrast.

After measuring the initial state of cells, we aspirated the medium and gently added 10 µM peptide in HBSS medium instead. We captured cells immediately after peptide addition and at 5, 10, 15, 20, 30, and 60 min after peptide addition.

Fiji/ImageJ was used for image analysis. We captured each confocal channel as a 3x3 montage and a 3-layer z-stack. First, among these z-slices, the best Tamra channel image was selected, along with the corresponding Hoechst channel z-slice. We cut this montage of 3x3 images and transformed it into a stack of 9 images, which were then background-corrected using the BaSic plugin [51] and subsequently stitched back into a single large image. Next, we determined the cell-nuclei area by transforming the Hoechst channel signal into a binary image. For each cell, the average Tamra channel fluorescence intensity signal in its nuclear area (defined in the Hoechst channel in the previous step) was calculated. Finally, this nuclear mean fluorescence value was compared with the mean Tamra channel fluorescence intensity in the outer cell space. If the nuclear fluorescence signal exceeded the background signal, the cell was considered penetrated. The weighted average of the percentage of cells penetrated, based on at least 3 independent replicates, is shown in Fig. 4C.

### Flow cytometry concentration study

U-2 OS cells cultured in 150 mm plates were trypsinised, resuspended in DMEM, and centrifuged for 5 min at 160 RCF. After aspiration of the supernatant, HBSS buffer was added. 1 ml of cell suspension was transferred into 1.5 ml Eppendorf tubes, and five samples were prepared: a cell-only control; 1 µM, 5 µM, 10 µM, and 15 µM Tamra-labelled Poli, Cmil, Human, or R_9_ peptides. These were incubated for 20 min, then centrifuged for 5 min in a mini centrifuge (Labnet Prism). The supernatant was aspirated, and cells in microtubes (1.5 ml) were put on ice to be promptly resuspended in HBSS for measurements performed on the CytoFLEX Flow Cytometer (Beckman Coulter). Measurements were done at room temperature, with low-speed acquisition and 10k target single cells. Poli, Cmil, and Human experiments were done in triplicate. A single replicate of R_9_ and R_4_ measurements was obtained from [40] for comparison. Singlet cell events were separated, and the fluorescence intensity histogram was obtained for this population, as shown in Supporting Fig. S3.

### Time-resolved flow cytometry

U-2 OS cells, detached from the cell culture dish by Trypsin treatment, were centrifuged and resuspended in the room temperature HBSS +/+ buffer. Immediately after the addition of 1 µM Tamra-labelled peptide to the cell suspension (10^5^ cells/ml), the sample was placed into the BD FACSAria™ Fusion flow cytometer (BD Biosciences). Tamra fluorescence was recorded in the PE-A channel (586/15 nm, YellGreen Laser 561) at an acquisition rate of 100 events/s (10 ms acquisition speed) with a 100-µm nozzle at room temperature. The time-dependent change in cell-associated fluorescence was used to monitor peptide association with the cells. The gating strategy used to identify the cell population for analysis is shown in Supplementary Fig. S4. Cells without peptide addition were used as a control and for calibration. Events were divided into three populations: singlet cells, aggregates, and subcellular events. For each of these populations, PE-A fluorescence intensity during measurement was recorded, as shown in Supporting Fig. S4. Supporting Fig. S5 confirms that peptides do not aggregate in HBSS +/+ buffer.

## Acknowledgements

KB, SJP and JZ thank the CB2 group for valuable discussions and support. KB acknowledges support from Charles University, where she is enrolled as a PhD student, and HPCg at IOCB Prague for computational resources. We thank P. Jungwirth for providing the peptides for measurements. We thank P. Jurkiewicz from the Imaging Methods Core Facility at BIOCEV, an institution supported by the MEYS CR (LM2023050 Czech-BioImaging), for his support with live cell imaging.

The research was supported by the Charles University grant Primus PRIMUS/23/MED/002, and by the Ministry of Health of the Czech Republic in cooperation with the Czech Health Research Council within the National Institute of Virology and Bacteriology, project No. NW26A-NIVB.

