## Supplementary material file for "The C-terminal region of interferon-γ mediates evolutionarily tunable membrane interaction"

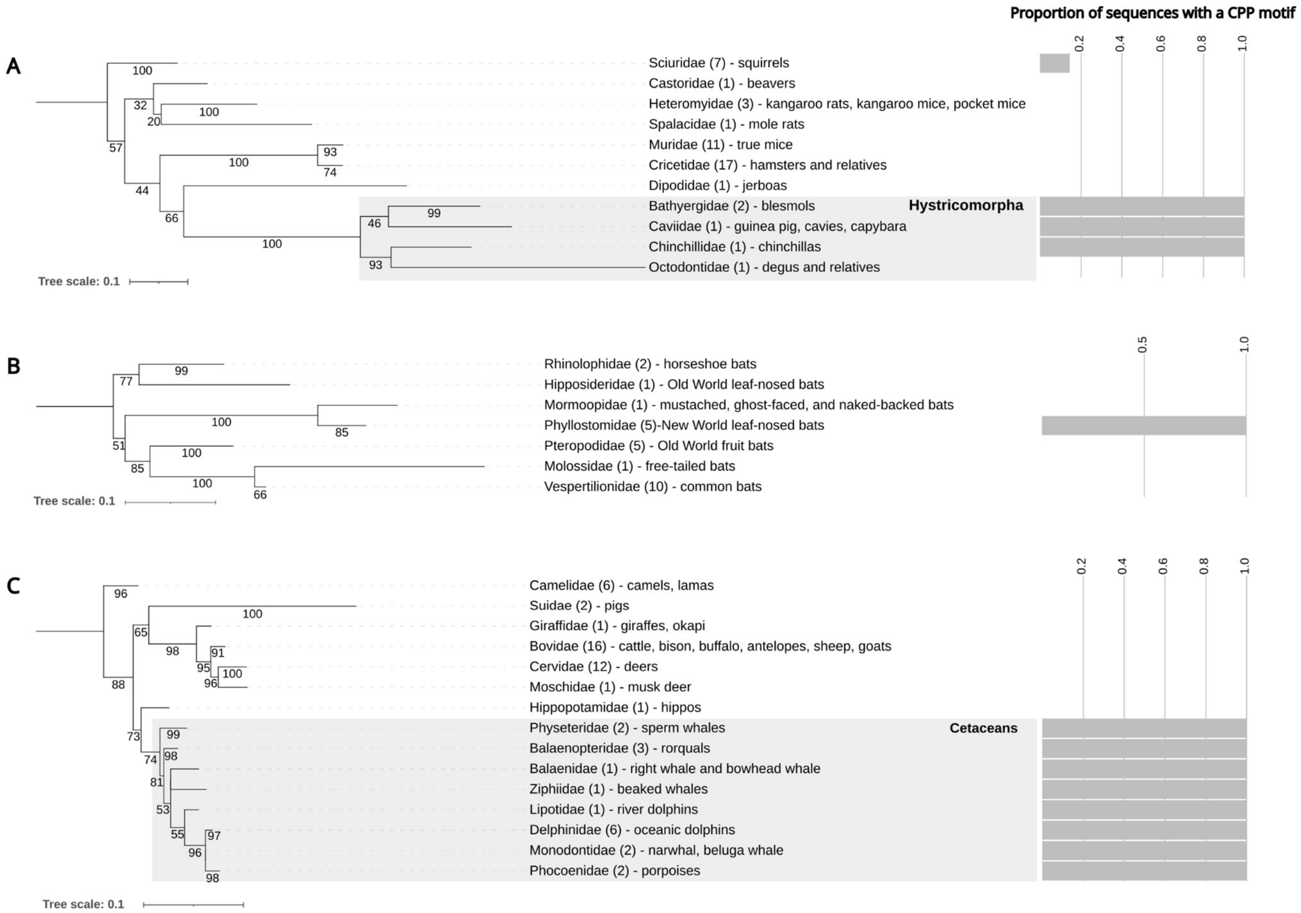


**Figure S1 – Presence of CPP motifs in mammalian species.** Maximum-likelihood phylogenetic trees (supported by 1000 bootstrap replicates) showing the evolution of IFNγ sequences in (A) rodents, (B) bats, and (C) artiodactyls. To the right, the bar charts depict the proportion of IFNγ sequences with a CPP motif in each family. All identified CPP motifs are the sequence of three arginines (RRR). The number in parentheses following the family name indicates the total number of IFNγ sequences identified in a given family. Bootstrap support values are shown at the corresponding nodes.

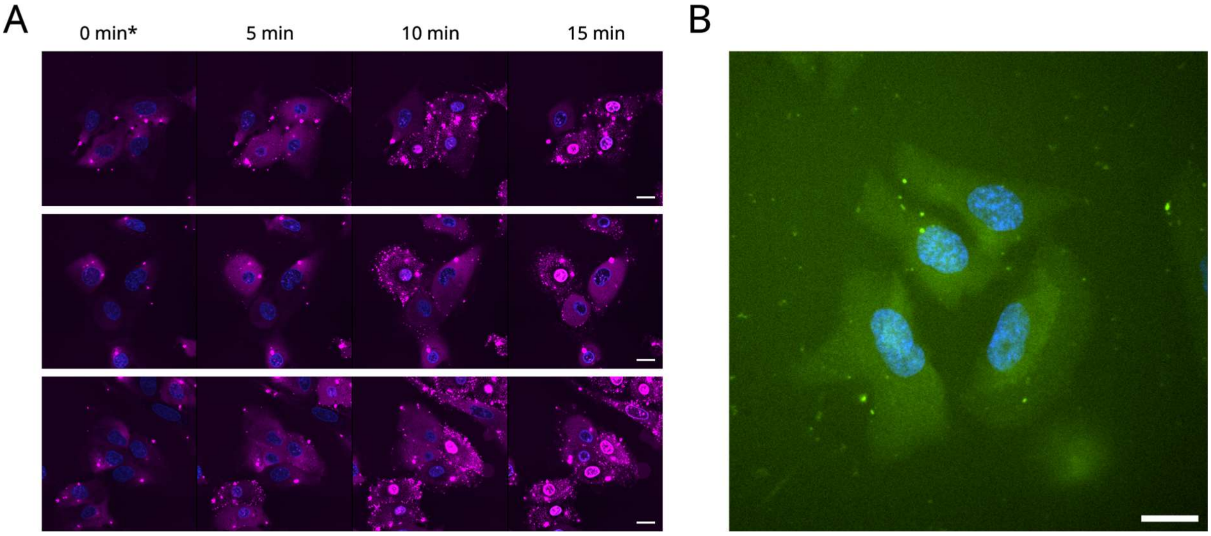


**Figure S2 – The effect of Tamra-labelled Poli peptide on U-2 OS cells.** (A) Live-cell imaging examples were measured immediately after adding 10 µM Tamra-labelled Poli (in red, denoted as 0 min*), and after 5, 10, and 15 min of incubation. Cell nuclei are labelled with Hoechst (blue). The scale bar is 20 µm. (B) UnaG green fluorescent protein fused with the Poli C-terminus sequence after 60 min U-2 OS cell incubation. The scale bar is 20 µm.

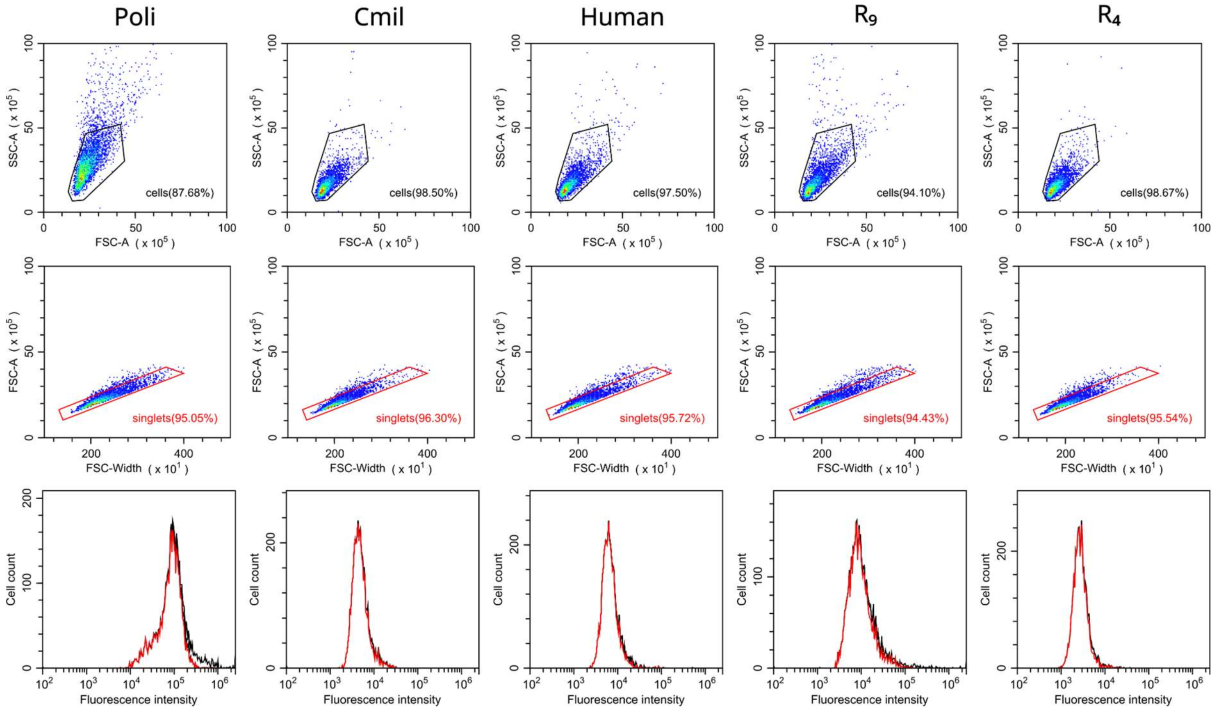


**Figure S3 – Flow cytometry gating strategy example.** We incubated U-2 OS cells with 10 μM Tamra-labelled peptides for 20 min, washed them, and examined them by flow cytometry. In the first step of the gating strategy (1st row), we selected viable cells. The second row shows singlet selection. Finally, a fluorescence histogram for the singlet subpopulation is shown in the third row. For comparison, the columns show peptides from left to right: Poli, Cmil, Human, $R_9$, and $R_4$. Note: $R_9$ and $R_4$ measurements were borrowed from Baxova et al. (2026) [40]

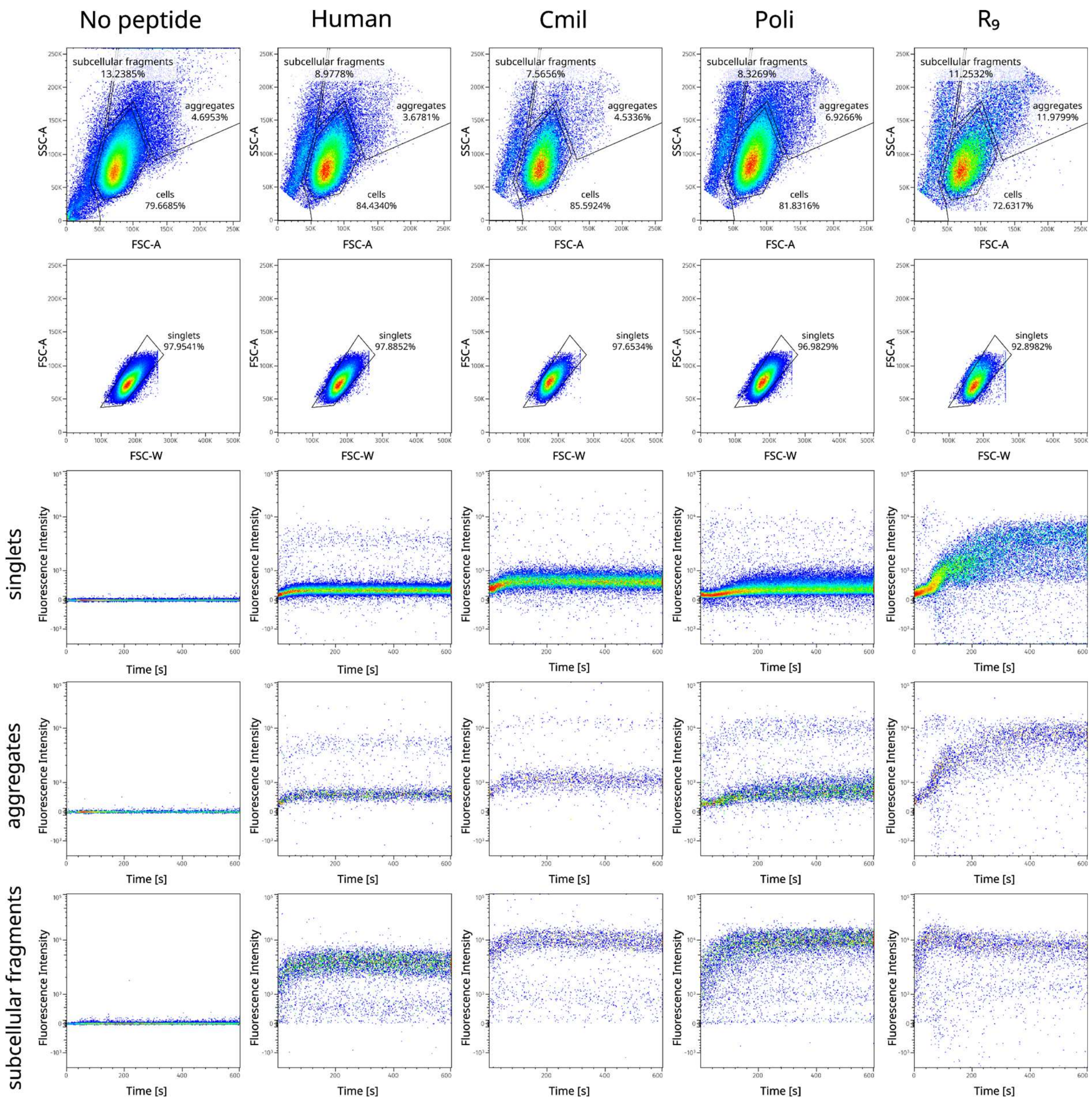


**Figure S4 – Time-resolved flow cytometry gating and results.** In the first step, shown in the 1st row, three cell populations are outlined: aggregates, subcellular fragments, and viable cell events. In the second step (2nd row), singlet cell selection is performed. Finally, fluorescence signal over time for singlets (3rd row), aggregates (4th row), and subcellular fragments (5th row) is obtained. The first column shows a control cell sample without any peptide addition. The next columns show Human, Cmi, Poli, and $R_9$ peptide measurements (from left to right). Note: $R_9$ measurement was borrowed from Baxova et al. (2026) [40]

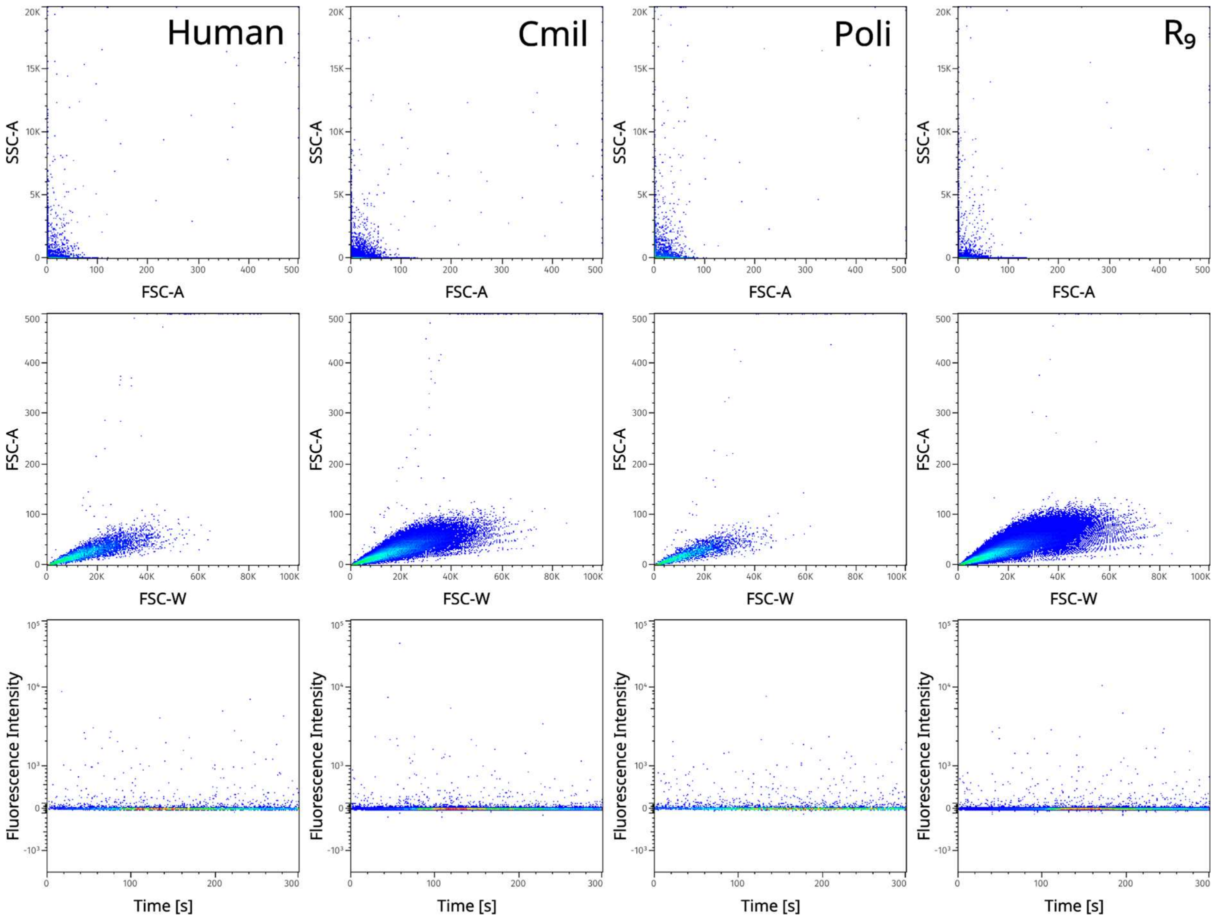


**Figure S5 – Checking for peptide aggregation by time-resolved flow cytometry.** In the absence of cells, we measured peptides in HBSS buffer by time-resolved flow cytometry. No prominent peptide aggregation occurs.

| Peptide | | | Poli | Cmil | Human | $R_9$ | $R_4$ |
|---|---|---|---|---|---|---|---|
| Peptide concentration | 0 µm | mean | 1532 ± 25 | 1464 ± 118 | 1549 ± 242 | 1773 | 2640 |
| | | median | 1390 ± 34 | 1310 ± 89 | 1383 ± 190 | 1545 | 1859 |
| | 1 µm | mean | 6187 ± 960 | 1865 ± 262 | 2034 ± 225 | 4049 | 1994 |
| | | median | 4278 ± 382 | 1649 ± 184 | 1816 ± 187 | 3416 | 1654 |
| | 5 µm | mean | 45545 ± 10296 | 4271 ± 688 | 4833 ± 730 | 8331 | 2335 |
| | | median | 33443 ± 10103 | 3468 ± 473 | 4018 ± 546 | 6062 | 1972 |
| | 10 µm | mean | 102719 ± 20715 | 6122 ± 919 | 8051 ± 164 | 11996 | 3114 |
| | | median | 91635 ± 15782 | 4822 ± 484 | 5885 ± 336 | 8518 | 2619 |
| | 15 µm | mean | 141735 ± 22907 | 7886 ± 1510 | 11896 ± 899 | 15007 | 3583 |
| | | median | 129264 ± 17792 | 6161 ± 958 | 8467 ± 1068 | 10536 | 2991 |

**Table S1 – Concentration-dependent flow cytometry results.** Fluorescence intensity distribution mean and median values obtained for singlet populations of U-2 OS cells incubated for 20 min with Tamra-labelled peptides of various concentrations.
